# Resolving Oligomeric States of Photoactivatable Proteins in Live Cells via Photon Counting Histogram Analysis

**DOI:** 10.1101/2025.04.18.649599

**Authors:** Tyler Camp, Zixiao Li, Yushan Li, Teak-Jung Oh, Kai Zhang

## Abstract

Oligomerization of photoactivatable proteins is widely used in optogenetics to modulate protein activity and regulate biological processes. However, their oligomerization states remain challenging to quantify in living cells. We applied photon counting histogram (PCH) analysis to quantify the oligomerization of two commonly used photoactivatable proteins, *Vaucheria frigida* Aureochrome light-oxygen-voltage (VfAuLOV) and *Arabidopsis Thaliana* cryptochrome 2 (AtCRY2), under dark and light conditions, providing a direct measurement of oligomerization states in live cells. Under blue light stimulation, VfAuLOV primarily forms dimers, whereas AtCRY2 forms higher order multimers in HEK293T cells. At intracellular concentrations above 1000 nM, AtCRY2 transitions into tetramers upon light stimulation, consistent with structural data obtained from cryo-electron microscopy. Unexpectedly, AtCRY2 (W374), a constitutively active mutant, still exhibits light-induced oligomerization. Human cryptochrome 2, in contrast, shows light-independent oligomerization at the intracellular concentration above 100 nM, below which the monomeric state dominates. Optogenetic signaling outcomes, demonstrated by light-induced lytic cell death, align with the oligomerization state of tested proteins. This study presents a quantitative framework for elucidating protein oligomerization in live cells, thereby enhancing the understanding of optogenetic mechanisms and their role in cellular signaling.

**Significance Statement:** Photoactivatable proteins enable the development of optical actuators that modulate cell signaling through light-mediated protein-protein interaction. Light-oxygen-voltage (LOV) and cryptochrome 2 (CRY2) are commonly used photoactivable protein modules that undergo blue light-mediated oligomerization, and the latter has been used to create synthetic biomolecular condensates in live cells. Although *in vitro* structural studies suggested their oligomerization states, their behavior in live cells is unclear. Here, we applied photon counting histogram analysis to determine the dark and light-activated oligomerization states of these photoactivatable proteins in live cells. Insights gained will provide better guidance in designing optical actuators, leading to more quantitative interpretation of the light-mediated signaling outcomes in cells.

## Introduction

Biomolecular condensates play a crucial role in catalyzing chemical reactions in live cells. Besides well-established intrinsically disordered proteins that can initiate the assembly of the condensates(1), photoactivatable proteins were used to modulate phase transition in live cells through light-induced protein-protein interactions, an optogenetic approach (2). Dimerization and oligomerization are fundamental principles in optogenetic system design (3). While in vitro methods, such as biochemical assays, crystallography, and Cryo-electron microscopy (Cryo-EM), provide insights into the oligomerization state of photoactivable proteins, a quantitative understanding of these dynamics in live cells is currently lacking. Such insights prove critical for rational design and practical implementation of optogenetic strategies in multicellular organisms, motivating a reliable, quantitative approach to resolve the *in situ* protein oligomerization state in live cells.

Fluorescence fluctuation spectroscopy (FFS), which relies on temporal fluctuations in fluorescence intensity, recovers information on fluorophore concentration and average oligomer size in live cells. Techniques falling under the FFS umbrella have been applied to study protein diffusion and oligomerization (4, 5) and include imaging techniques such as spatial intensity distribution analysis (SpIDA) (6) and fluorescence intensity fluctuation (FIF) analysis (7). FFS-based experiments are particularly useful for studying assemblies larger than dimers that are challenging to resolve by FRET (8). Photon counting histogram (PCH) analysis (9) and related single-point techniques, wherein the fluorescence signal is measured at a single spot over time using focused, stationary laser excitation, has successfully resolved dimers (10), pentamers (11), and even much larger assemblies of more than several hundred protein subunits per complex (12), all within live cells. Here, we use PCH analysis to resolve the oligomerization state of photoactivatable proteins in live cells. We found that the HaloTag, a self-labeling protein tag, serves as a better probe for PCH analysis than conventionally used fluorescence proteins (13), due to improved brightness and photostability of chemically synthesized small-molecule fluorescent probes.

To gain insights into light-mediated dimerization/oligomerization of photoactivatable proteins in live cells, we selected two commonly used homotypic dimerization/oligomerization photoactivatable proteins: the light-oxygen-voltage (LOV) domain of *V. frigida* Aurochrome1 (herein VfAuLOV) (13–15) and the photolyase homology region of cryptochrome 2 from *A. thaliana* (herein AtCRY2) (16). Additional derivatives of these proteins, bZip-VfAuLOV (17) and the constitutively active AtCRY2 W374A mutant, as well as the human CRY2 photolyase homology region (HsCRY2), were also tested. Previous structural studies used size-exclusion chromatography to resolve aggregation in vitro (17), together with the crystal structures of AtCRY2 resolved by Cryo-EM (18–20) or X-ray crystallography (21, 22). However, only low-resolution descriptions of photo-induced protein-membrane associations (23–25) or homo-hetero-oligomerization (26–29) were available in cells. Measuring the oligomerization state of these optogenetic systems in live cells would therefore fill an important gap and inform future cellular engineering efforts toward a mechanistic understanding of the relationship between protein concentration, oligomerization state, and signaling outcomes.

## Results

### Fluorescent proteins show distinct photostability in mammalian cells

Conventional FFS often uses fluorescent proteins (FPs) as the fluorophore. An optimal FP should have high brightness (proportional to the product of absorption coefficient and quantum yield) and photostability. Thus, we first compared the photostability of various commonly used fluorescent proteins (FPs), including mEGFP, mGold, mCherry, and mScarlet. HEK293T cells were transfected with each plasmid and incubated for 16 hr before imaging. A small region was selected from each cell to analyze photobleaching data, and the average intensity was measured for each frame. Representative photobleaching trajectories of each FP are shown in **Figure 1A**. The photobleaching rate constant for each FP was calculated by fitting each curve with a single-exponential decay. To calibrate the effect of distinct excitation wavelengths, we plot the recovered photobleaching rate constants against E_c_, which measures the amount of light power coupled into the chromophore of each FP (**Figure 1B** and Materials and Methods). Results showed that mEGFP and mCherry outperformed (i.e., showed smaller photobleaching rate constants than) mGold and mScarlet. This finding is consistent with the literature, which shows the reduced stability of mScarlet relative to mCherry (29). However, the low stability of mGold was surprising, given previously published results (30).

**Figure 1.**
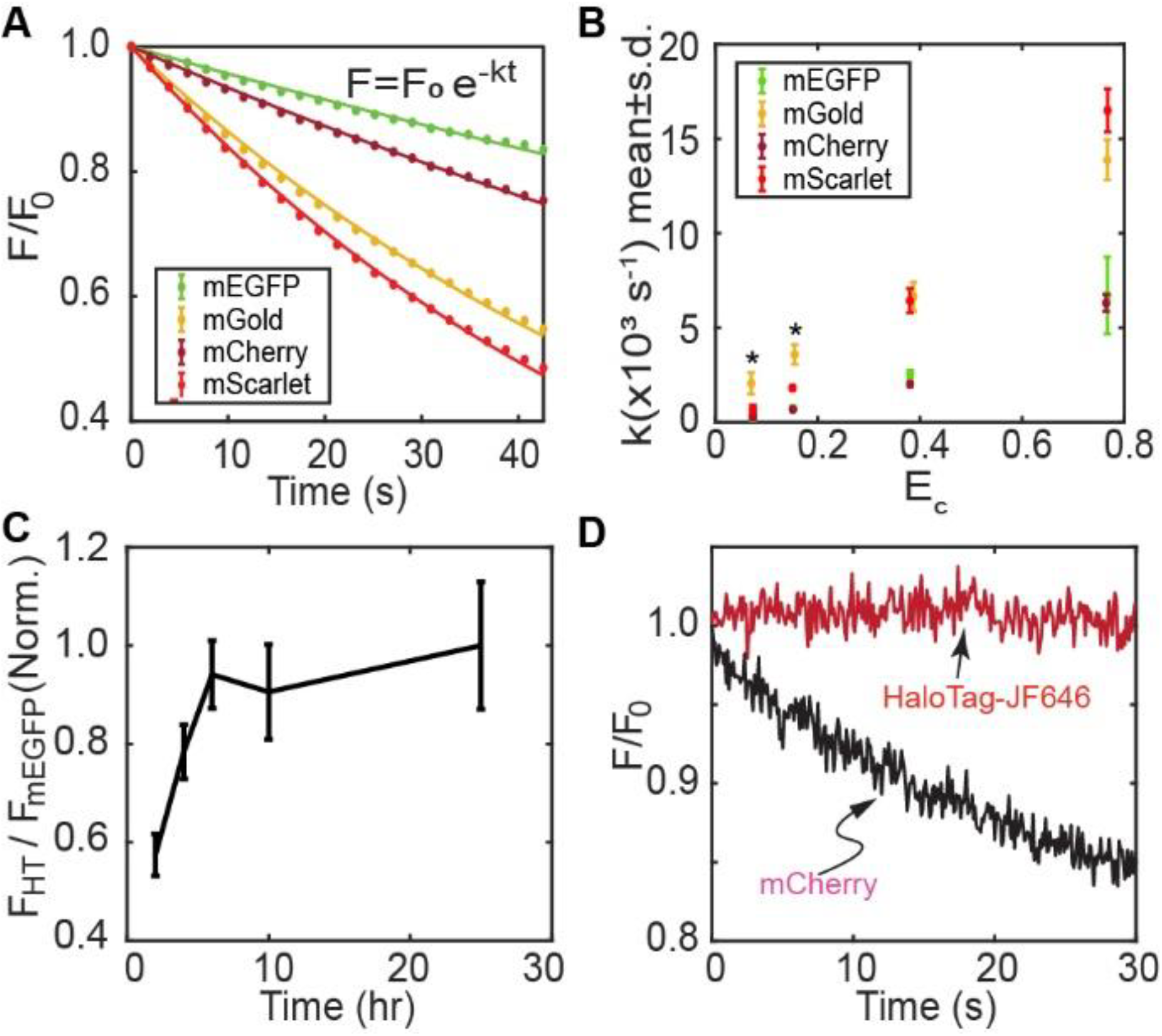
Benchmark fluorophore photophysics in live cells. (A) Representative photobleaching trajectories from cells expressing mEGFP, mGold, mCherry, and mScarlet after repeated scanning on a commercial laser scanning confocal microscope. (B) Photobleaching rate constants recovered from the bleaching of individual cells as a function of relative excitation power. Asterisks mark the apparent rate constants for mGold excited at low power, where a single-exponential model did not fully describe the fluorescence decay. (C) Optimization of HaloTag-JF646 staining protocol. Cells expressing a fusion of HaloTag and mEGFP were stained with JF646 for the indicated times, destained, and imaged using a plate-based fluorescence imaging system. (D) Single-spot photobleaching trajectories from cells expressing HaloTag-JF646 (red) or mCherry (black) that were excited at a similar power. (A) and (D) Data is normalized to the intensity at t = 0.

### HaloTag-JF646 outperforms FPs in photostability

The HaloTag protein is a modified haloalkane dehalogenase that has been engineered to covalently bind fluorescent ligands with exceptional photostability and brightness (31). Hence, they are excellent candidates for use in FFS-based measurements. However, many quantitative imaging studies continue to use FPs despite their low photostability and the presence of blinking, dark, or low fluorescence states (32). In the context of FFS experiments, we surmised that HaloTag might outperform traditional FPs based on the high brightness and photostability of HaloTag ligands (33). To optimize the HaloTag labeling procedure, we constructed a tandem HaloTag-mEGFP plasmid. Complete labeling of HaloTag should result in a stable fluorescence ratio of labeled HaloTag to mEGFP. Complete HaloTag labeling was achieved after 6 hours of incubation of 200 nM JF646-HaloTag ligand (**Figure 1C**). Optimization of HaloTag labeling is crucial for accurately interpreting quantitative measurements of protein oligomerization, as incomplete labeling can lead to a severe underestimation of the actual oligomerization state(s) in PCH analysis. We then compared the photostability of HaloTag-JF646 and mCherry. Under similar excitation power, JF646 remains stable, while 15% mCherry was photobleached within 30 seconds of illumination in HEK293T cells (**Figure 1D**).

### PCH resolves molecular brightness in live cells

To confirm that PCH analysis could resolve the brightness of a monomer from a constitutive dimer, we collected fluorescence fluctuation trajectories in HEK293T cells expressing either monomeric or tandem dimeric mEGFP, mCherry, and HaloTag (**Figure 2A-B**). To calculate the molecular brightness for each construct, we employed PCH analysis with a 20 kHz sampling frequency and a detector dead time of 15 ns (see Materials and Methods). Although a dimer should have a molecular brightness twice that of a monomer, various factors, such as protein misfolding, chromophore maturation, and the presence of dark or low-fluorescent states of the chromophore, could result in a lower apparent brightness for the dimer (34). As an example, monomeric EGFP is known to occupy a fluorescent state with a probability of 80% (35). To compute *p*_*f*_, the probability that each fluorophore is fluorescent, we apply the following model (34): 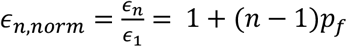, where the subscripts represent molecular brightness values of the monomer or an *n*-mer. We use the experimentally determined ratio of the molecular brightness of the dimer and monomer (ϵ_2_/ϵ_1_) to calculate the probability that each fluorophore is fluorescent. Results showed that the dimer:monomer ratio of the median brightness values (ϵ_2_/ϵ_1_) for mEGFP and mCherry are 1.52 and 1.32, respectively, which is close to but lower than previously reported values of 1.63 and 1.42 recovered from brightness imaging (34). The small differences might result from the use of a different linker sequence that separates the protomers in the tandem dimer constructs, which may affect the folding efficiency of the dimers. In contrast, the dimer:monomer brightness ratio of HaloTag-JF646 was 1.70, indicating that more HaloTag protein molecules are fluorescent relative to mCherry (**Figure 2C**). Besides enhanced photostability, the redshifted absorption peak of JF646 allows it to be used as an “orthogonal” label that can be observed without simultaneously activating the commonly used optogenetic proteins. For example, both VfAuLOV and AtCRY2 can be activated by 405-488 nm light, overlapping with the absorption spectrum of mEGFP.

**Figure 2.**
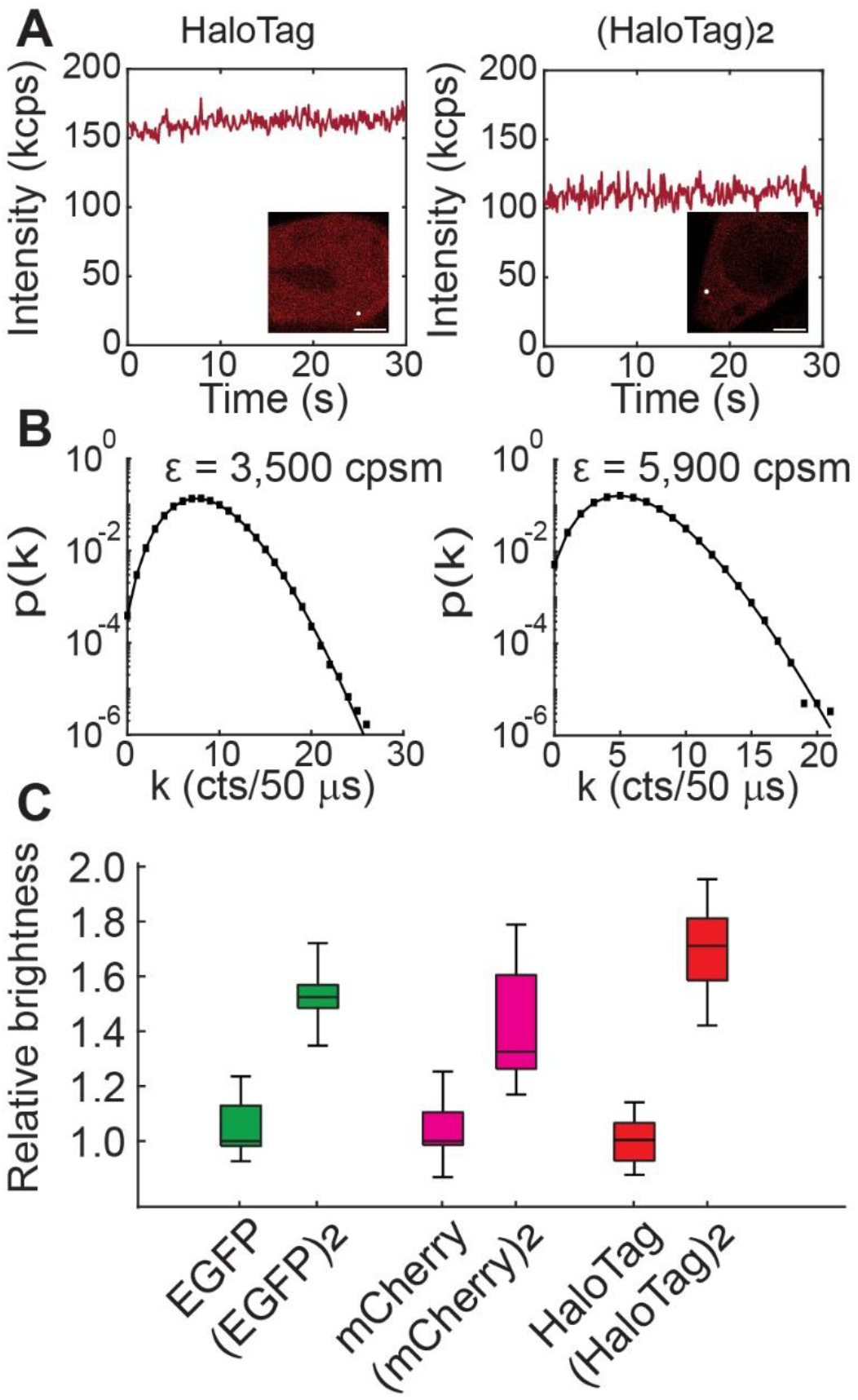
HaloTag-JF646 outperforms fluorescent proteins for quantifying protein oligomerization in live cells. (A) HEK293T cells expressing a HaloTag monomer (left) or tandem dimer (right) stained with JF646 were imaged using a custom confocal system, and individual spots were selected for FFS analysis (white spot in inset image, scale bars: 6 µm). (B) Fluorescence fluctuations were fit to the PCH model to retrieve brightness in units of counts per second per molecule (cpsm). Black squares represent the data while the black line shows the fit to the PCH model. (C) The relative molecular brightness of each labeling system.

### PCH confirms blue light-induced dimerization of VfAuLOV in live cells

Next, we applied PCH analysis to the VfAuLOV system. Quantification of VfAuLOV’s oligomerization state has so far relied on reconstituted proteins in vitro (17). It remains unclear whether fusion proteins containing VfAuLOV form primarily dimers or higher-order oligomers in cells in response to blue light stimulation. To quantitatively determine the light-induced oligomerization of VfAuLOV, we transfected HEK293T cells with HaloTag-VfAuLOV and then stained them for 6 hours with JF646 HaloTag ligand (**Figure 3A**). We first measured the molecular brightness of a population of cells in the dark state. These cells exhibited a brightness comparable to that of the monomer control. By converting the fitted number density recovered by PCH into concentration using a calibration measurement (**Figure S1** and Materials and Methods), we determined that the intracellular protomer concentration ranges from 50 nM to 1200 nM (**Figure 3B**, black circles). Next, we applied 5 minutes of continuous blue light stimulation (6 mW/cm^2^ measured at 488 nm) to the same cells on the microscope. We normalized the relative brightness to account for incomplete labeling based on the observed HaloTag dimer:monomer brightness ratio of 1.70 (34). This correction is applied to all relative brightness values reported for our optogenetic constructs. After blue light exposure, the relative brightness approaches 2, consistent with the formation of dimers.

**Figure 3:**
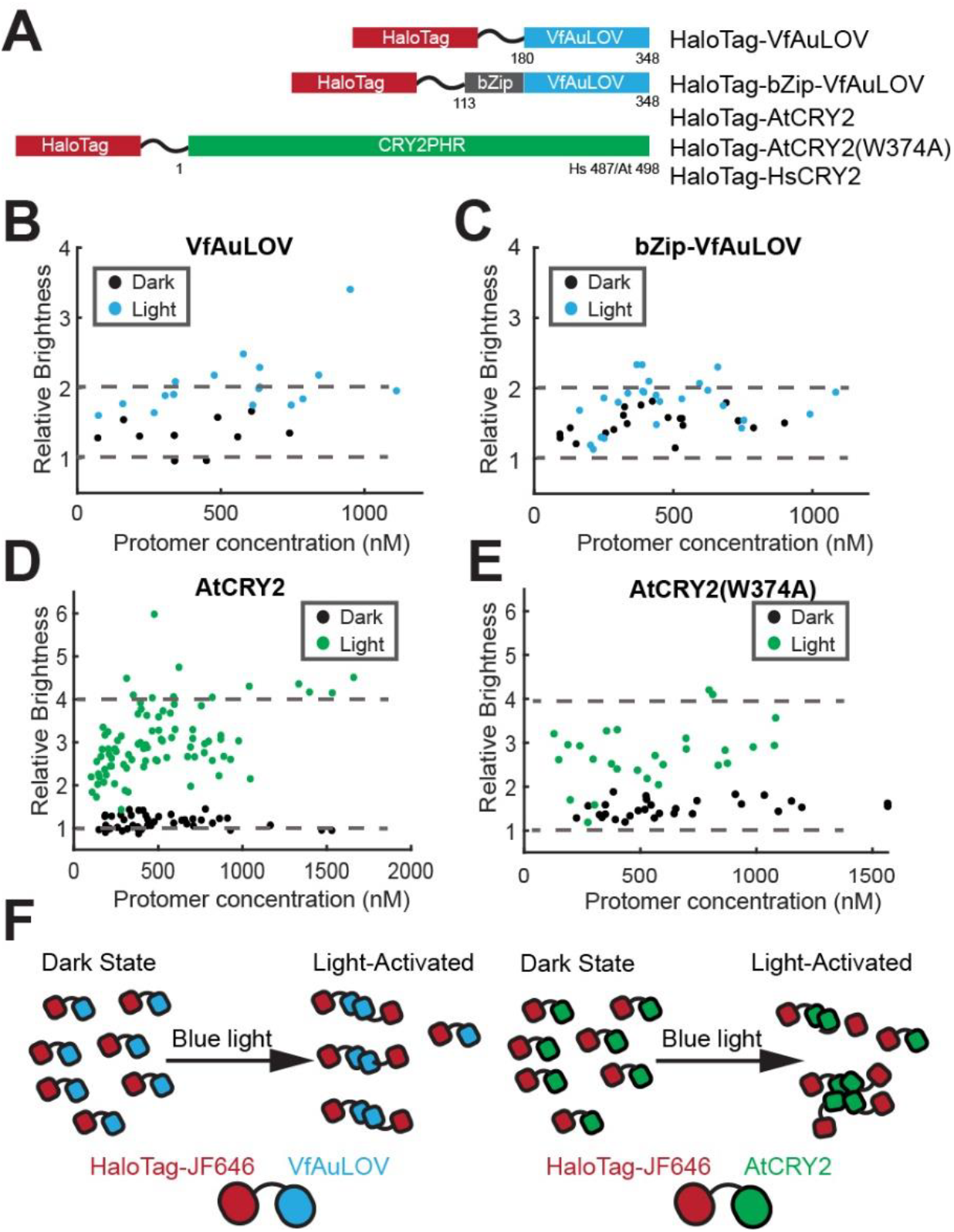
Oligomerization state of AuLOV and CRY2 in live cells. (A) Constructs used in this work. Residue numbering is shown according to the numbering of the full-length proteins. (B) Relative molecular brightness of VfAuLOV (wild-type) in the dark (black) and light-activated (blue) states. Brightness values are relative to the monomeric standard. (C) Same as (B) but for bZip-VfAuLOV. (D-E) same as (B) but for wild-type AtCRY2 and the constitutively active W374A mutant. (F) PCH-inspired model for photoactivatable proteins in live cells. In the dark state, VfAuLOV and AtCRY2 exist primarily as monomers. After blue light stimulation, the equilibrium shifts toward a dimeric (VfAuLOV) and tetrameric (AtCRY2) conformation. (B-E) Dotted lines are drawn at the expected brightness of a fully monomeric or dimeric (VfAuLOV constructs) or tetrameric (AtCRY2 constructs) population.

For light-exposed cells, the HaloTag fusion constructs are expected to exist as a population of mixed oligomers. For this reason, the brightness values (ε) and number of particles (N) reflect weighted averages of the mixed population. For example, VfAuLOV is likely to exist as a mixture of monomers and dimers after light exposure, with the dimer fraction depending on the self-dissociation constant and total protein concentration. The “apparent” or weighted-average molecular brightness is given by the following equation (9) 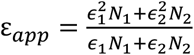. The fitted value of N_*app*_ reflects not the number of individual protein molecules but the “apparent” number of molecules that correspond to the average brightness value of ε_*app*_. To calculate the true protein concentration, we note that the following relation holds for all samples: ε_*app*_N_*app*_ = ε_1_N_1_, where the right-hand terms ε_1_ and N_1_ are the monomeric brightness and the number of monomer equivalents (or protomers), respectively.

Occasionally, we measure higher relative brightness values (>3) that could indicate higher-order aggregates. Alternatively, higher brightness values could arise from cell movement during the measurement, which could periodically shift the observation volume into a different area with a different local concentration, artificially increasing the observed intensity fluctuations. When basic residues and a leucine zipper domain are included (bZip-VfAuLOV), a base-level oligomerization was shown in the dark, which we attribute to the stabilizing effect of the basic/zipper region. Like VfAuLOV, bZip-VfAuLOV displays light-induced dimerization (**Figure 3C**). Thus, both constructs dimerize when exposed to blue light in live cells, with the bZip extension resulting in higher basal oligomerization in the dark.

### AtCRY2 forms multimers upon blue light illumination in live cells

We then measured the light-dependent oligomerization of AtCRY2 in HEK293T cells expressing HaloTag-AtCRY2 (**Figure 3A, 3D-F**). The dark state brightness was consistently shown to be monomeric in cells expressing up to 1600 nM of protein (**Figure 3D, black circles**). After exposure to blue light, the relative brightness values increase to mostly between 2 and 4 (**Figure 3D, green circles**), indicating that AtCRY2 exists as mixed oligomers when exposed to blue light. Near-full conversion of AtCRY2 into tetramers was observed in cells expressing more than 1,000 nM proteins. Next, we applied PCH to study AtCRY2(W374A), a constitutively active mutant in plants, even in darkness (36). This mutant has been observed to form dimers and tetramers *in vitro* (18). We observe a relative brightness of about 1.5 in the dark, indicating less than full conversion to either dimers or tetramers (**Fig 3E, black dots**). Unexpectedly, the W374A mutant still undergoes light-induced oligomerization (**Fig. 3E, green dots**). This is surprising given that the W374A mutation severely reduces photoreduction *in vitro* (36). However, the authors of the same study also noted that the W374A mutant still responds to changes in light fluency. Additionally, AtCRY1(W400F), which lacks an electron donor to FAD, exhibits decent photoreduction in insect cells when exposed to 150 µmol m^-2^ s^-1^ (approximately 4 mW/cm^2^) light at 450 nm, likely due to a more conducive redox environment compared to in vitro conditions (37). Thus, we conclude that the W374A mutant has a higher basal oligomerization state and retains the ability to oligomerize further upon light stimulation in cells. Taken together, these results reveal distinct photooligomerization behaviors between VfAuLOV and AtCRY2 (**Fig. 3F**).

### Human CRY2 shows concentration-dependent, light-independent oligomerization in live cells

Animal cryptochromes are primarily photoreceptors that perform both light-dependent and light-independent functions in the regulation of the circadian clock (38). For example, human cryptochromes are believed to be a light-independent transcription factor. However, evidence shows that human cryptochrome (HsCRY1) can respond to light and undergo photoreduction in cells. Previous co-immunoprecipitation experiments show light-independent oligomerization of HsCRY1 and HsCRY2 in HEK293 cells (29), but the stoichiometry of their conformational state is unclear. To define the oligomerization state of human cryptochrome in cells, we applied PCH analysis to HaloTag-HsCRY2. At low intracellular concentration (less than 100 nM), HsCRY2 appears to stay as monomers in the dark and after blue light stimulation. The average brightness increases at higher concentration, irrespective of light exposure, reaching 1.6 at about 750 nM, indicating a mixture of oligomerization states (**Figure 4A**). In comparison, AtCRY2 shows a higher oligomerization state with an average brightness of approximately 3 at 750 nM after blue light stimulation (**Figure 3D**).

**Figure 4:**
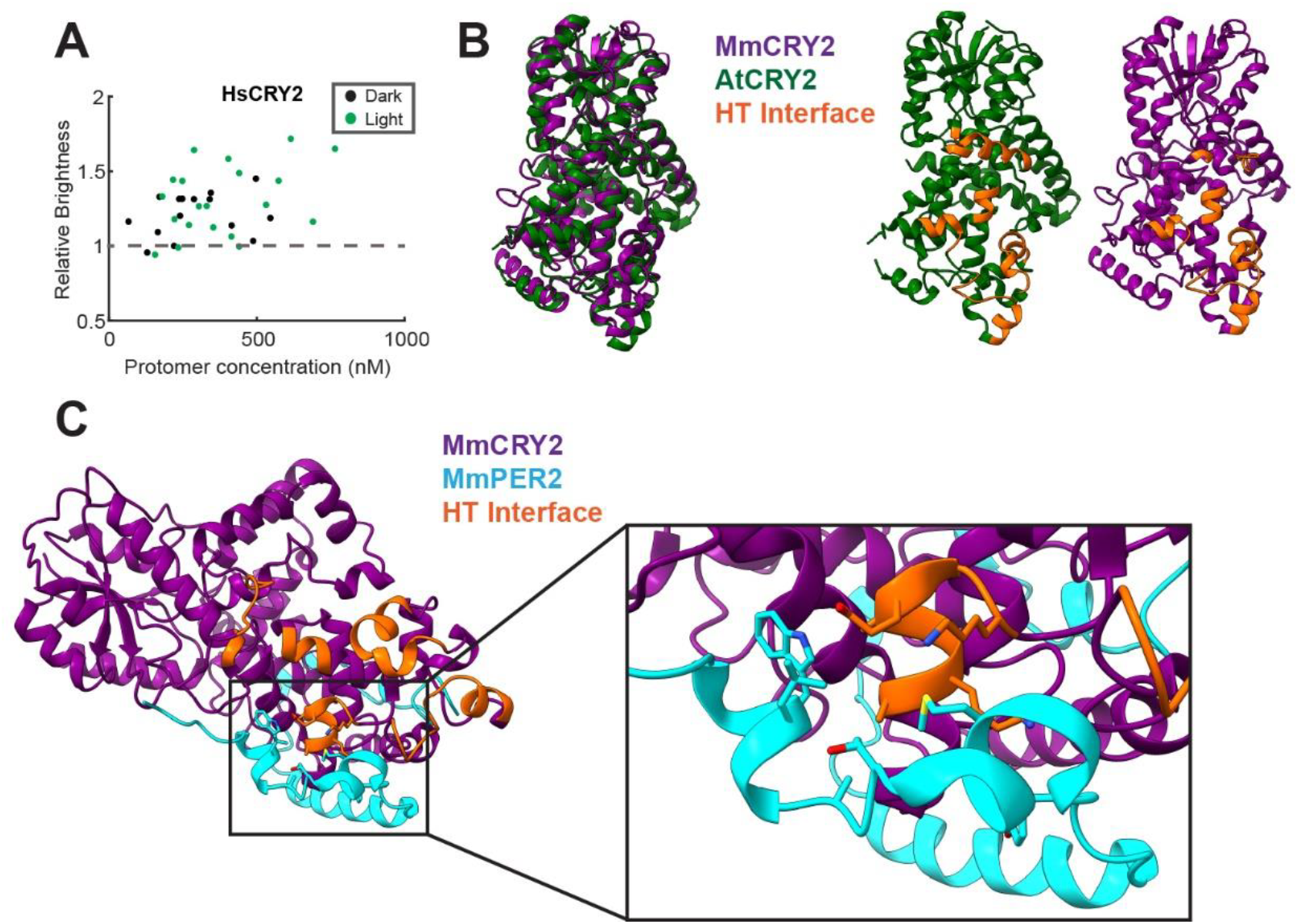
Occlusion of the HT interface by PER2 in mammalian CRY2. (A) Relative molecular brightness of HsCRY2 in the dark (black) and light-activated (green) states. Brightness values are relative to the monomeric standard. (B) (left) Structural alignment of AtCRY2 (PDB 6K8I) with MmCRY2 (PDB 4I6G) showing conserved overall fold of the PHR domains. (right) Separate renderings of MmCRY2 (purple, right) and AtCRY2 (green, left) showing the residues involved in the HT interface in AtCRY2 (orange). (C) Crystal structure (PDB 4U8H) of MmCRY2 (purple) in complex with the CRY-binding domain of murine PER2 (cyan) showing partial occlusion of the putative HT interface (orange).

We wondered if the same residues involved in self-oligomerization in AtCRY2 might be involved in hetero-oligomerization with other effector molecules in mammals. We first aligned the published structure of FAD-bound *Mus musculus* CRY2 (MmCRY2, PDB 4I6G), which shares ∼95% amino acid sequence identity with HsCRY2, with the inactive AtCRY2 monomer (PDB 6K8I). This alignment showed a moderate RMSD (4 Å) but a similar overall fold (**Figure 4B**). The resulting structural alignment allowed us to identify the residues in MmCRY2 that form a putative Head-to-Tail (HT) interface (**Figure 4B, Figure S2**), which has been previously identified in the oligomeric structure of AtCRY2 (22). Next, we analyzed the published crystal structure of the CRY-binding domain of murine PER2 with MmCRY2 PHR (39) (PDB 4U8H) and found that the HT interface is partially occluded (**Figure 4C**). This analysis reveals that the same residues involved in self-oligomerization in AtCRY2 may have evolved to promote hetero-oligomer formation in mammals, consistent with a model of light-independent signaling of HsCRY2. Our data do not rule out the possibility that light influences the physical interactions of HsCRY2 with other effectors. Whether this is true requires direct experimental testing of light-dependent binding of HsCRY2 to effector molecules, such as PERs, which are known to compete with FAD for binding to MmCRY2 (40). It is also unknown whether FAD binding to HsCRY2 is influenced by light absorption of FAD, although FAD has been observed to stabilize CRY2 protein in cell culture and in mice (41).

### The potency of optogenetically induced lytic cell death through RIPK3 oligomerization correlates with the oligomerization state of VfAuLOV and AtCRY2

To link the oligomerization behavior of photoactivatable proteins with their signaling outcomes, we fused mCherry-RIPK3 (receptor-interaction protein kinase 3) to the N-termini of VfAuLOV, bZip-VfAuLOV, and AtCRY2. RIPK3 clustering leads to lytic cell death, which can be stained by SytoxGreen, a membrane-impermeable nucleic acid staining dye (**Figure 5A**). We reasoned that the death-inducing potential from each fusion construct differs due to their distinct oligomerization behavior in the dark and light. When expressed individually in cells, the basal level of cell death ranks as bZip-VfAuLOV > VfAuLOV > AtCRY2, which is consistent with their dark oligomerization states. Blue light-induced cell death potential ranks as AtCRY2>bZip-VfAuLOV∼VfAuLOV, suggesting that a higher oligomerization state of RIPK3 induces more potent cell death, consistent with previous studies (42).

**Figure 5.**
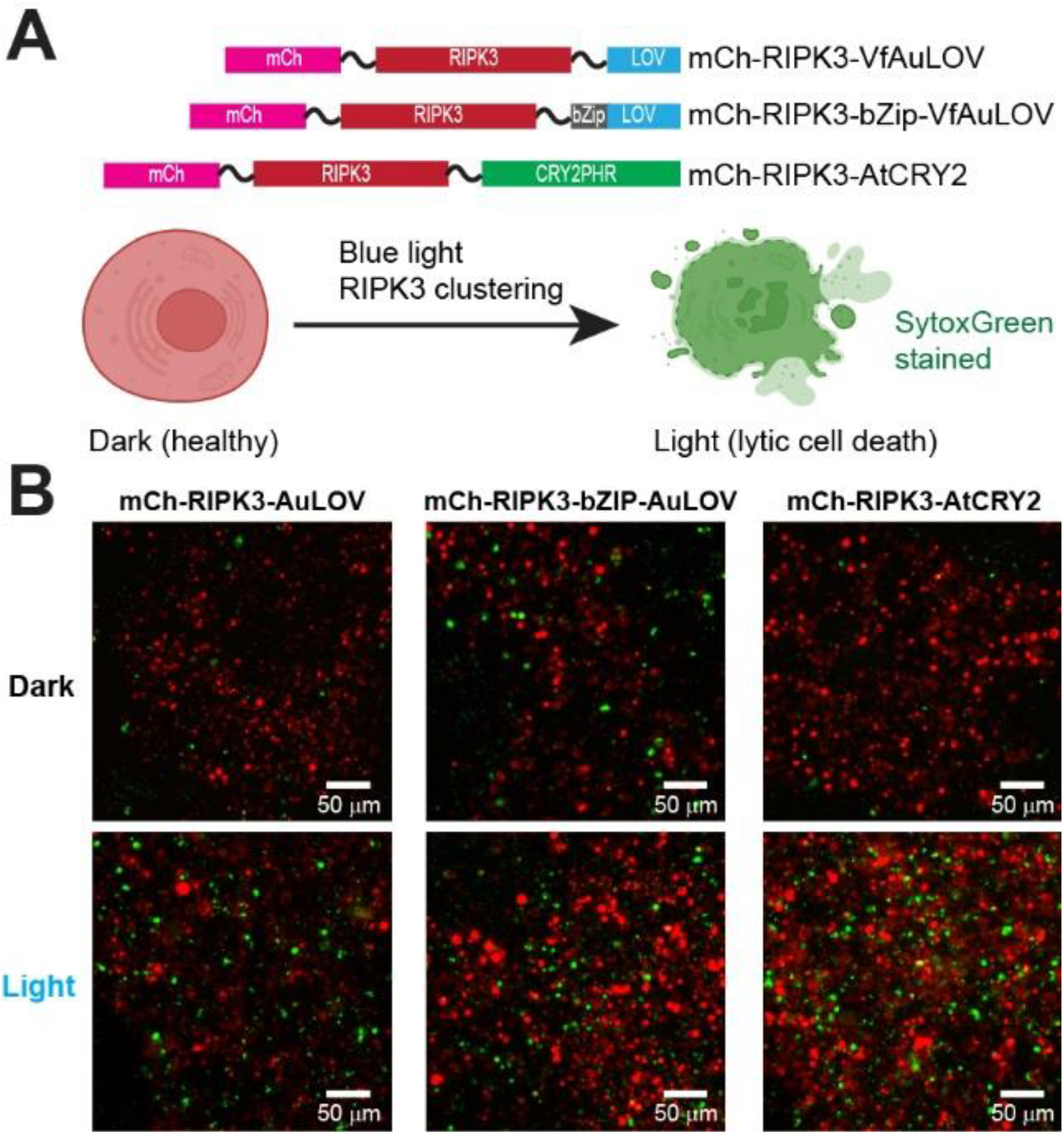
The potency of optogenetically induced lytic cell death depends on oligomerization states. (A) (Top) Constructs used in this work. (Bottom) Light-induced clustering of RIPK3 is expected to cause lytic cell death (necroptosis) marked by SytoxGreen staining. (B) HEK293T cells transfected with each construct under dark and light treatment.

## Discussion

This study characterizes the blue-light-induced homotypic protein association of VfAuLOV, AtCRY2, and HsCRY2, which are expressed in HEK293T cells at intracellular concentrations ranging from 50 to 1700 nM and illuminated with mild light power (6 mW/cm^2^). This light power is chosen because it falls within the range of typical blue light LED output, comparable to the physiological conditions on the sunlight-illuminated Earth’s surface. The blue spectrum of solar irradiance is estimated to be 14.3 mW/cm^2^, assuming 42% of the total solar irradiance (1361 W/m^2^, as measured by NASA) is visible light, and the blue spectrum (400-500 nm) accounts for 25% of the visible power. Additionally, this power (6 mW/cm^2^, equivalent to 225 µE m^-2^ s^-1^ or 225 µmol m^-2^ s^-1^ at 450 nm) has been validated by multiple research laboratories, which enables the robust stimulation of various LOV- and cryptochrome-based optogenetic responses in vitro (36, 43) or in cells (44, 16, 45–48, 23, 24, 29). The range of 50-1700 nM intracellular protein concentration corresponds to the average expression level of the Ubiquitin C (UBC) promoter, which has a weak promoter strength that measures about 10% of that of the cytomegalovirus (CMV) promoter (49). The rationale for selecting the UBC promoter was twofold: to minimize the side effects of overexpression and ensure that sufficient fluctuation can be measured in PCH analysis, as higher concentrations imply less relative fluctuation. Under these conditions, our data confirm that VfAuLOV predominantly forms dimers upon blue light illumination in live HEK293T cells, consistent with structural information gained from biochemical analysis (17).

This study provides new insights into the activation of AtCRY2. In cells, AtCRY2 forms monomers in the dark but transitions to tetramers at intracellular concentrations exceeding 1,000 nM under blue light stimulation, consistent with biochemical analysis (1) and the CryoEM structure of the tetrameric structure (19). A mixture of oligomer states (likely monomer, dimer, and tetramer) was observed in cells expressing 50-1000 nM, with an average relative brightness between 2 and 4. The average relative brightness is positively correlated with the concentration of protomers. However, AtCRY2(W374A), a constitutively active mutant with a dimeric and tetrameric structure resolved by CryoEM in the dark (18), shows only a slightly increased relative brightness (∼1.5) in PCH analysis up to 1600 nM, suggesting that active signaling tetrameric state formation may require even higher concentrations in cells.

Unexpectedly, W374A still exhibits light-induced oligomerization. Since photoreduction is severely compromised in the W374A mutant, the observed light-induced oligomerization supports the hypothesis that its oligomerization does at most partially require photoreduction via the Trp-triad (W321, W374, and W397) in AtCRY2. Interestingly, previous studies reveal that AtCRY2(W374A), despite being constitutively active, still exhibits activity-inhibiting effects on hypocotyl elongation in response to blue light for unknown reasons (36). Our PCH result suggests a possible explanation that such a response could result from blue light-induced W374A oligomerization. Inspired by the previous structural study (18), we speculate that blue light illumination excites the FAD cofactor, causing a small conformational change that propagates and amplifies larger conformational changes away from the FAD binding pocket. Such amplified conformational changes, likely near the N-terminus, facilitate CIB1 binding, as suggested by mutational studies (28) and CryoEM structures (20), and stabilize the tetrameric state of AtCRY2. Importantly, such a conformational change originating with FAD photoexcitation does not require electron donation from the Trp-triad.

Besides brightness, PCH analysis also directly measures the intracellular concentration of photoactivatable proteins, which may provide insights into the quantitative interpretation of their photo-oligomerization behaviors. For example, previous work suggests HsCRY2 could homo-associate in a light-independent manner (29, 50). Here, PCH analysis indicates that HsCRY2 primarily exists as monomers in darkness and light at 100 nM but shows slightly increased oligomerization as the concentration increases. Such a difference could be explained by the fact that a weaker promoter (UBC) was used in this work, in comparison with the stronger CMV promoter (approximately 10-fold stronger than UBC). Thus, we speculate that HsCRY2 could undergo homo-association at an intracellular concentration between 1000 nM and 12 µM (10-fold of UBC-driven protein expression) in HEK293T cells. Of course, oligomerization is only one potential conformational state for AtCRY2 signaling. A recent study reveals that AtCRY2 functions even in the dark through the binding to effector proteins of Forkedlike 1 (FL1) and FL3, and blue light inhibits such interactions (51). This further supports that AtCRY2 signaling depends more on its selective binding to effector proteins, either in darkness or light.

Technical advances of this work include a comparative analysis of organic HaloTag ligands and FPs. HaloTag-JF646 outperforms all FPs tested in this work (mEGFP, mCherry, mGold, and mScarlet) in terms of photostability and an improved dimer:monomer molecular brightness ratio compared to FPs, making it a more suitable candidate for PCH analysis. However, caution should be exercised to ensure complete HaloTag labeling. Based on our experience, labeling time is the most critical factor, and 6 hours is used for complete labeling of HaloTag in live cells. We emphasize that each chemical label, even from the same manufacturer, must be tested independently to ensure optimal labeling in the cell line used. We also noted limitations of the current PCH analysis, as it only measures population averages; it cannot directly resolve the fraction of monomers, dimers, tetramers, etc., from a mixed population. Additionally, our measurements following light exposure are collected for 30 seconds after the light is turned off, during which some of the complexes may dissociate (**Figure S3**). Future experiments to address this limitation, such as cell fixation during blue light exposure followed by super-resolution imaging, are ongoing in our laboratory.

## Materials and Methods

### Home-built Microscope Setup

We outfitted our Olympus IX73 microscope (Evident) with a galvanometer-driven scanning mirror unit provided by ISS Inc. (Champaign, IL). To observe transfected cells before FFS imaging, we coupled the laser excitation into the back port of the microscope and illuminated the sample dish in epifluorescence mode. The fluorescence was observed through the eyepiece, and cells of interest were centered by manually moving the translation stage. Alternatively, we used the built-in incandescent light of the IX73 to locate the focal plane containing cells before switching on the laser light for FFS measurements.

To augment the system with confocal scanning, the excitation lasers were split from the epifluorescence light path with a 30/70 beamsplitter (BSS10, Thorlabs). Our system includes three laser lines: 488 nm 100 mW c.w. and 561 nm 50 mW c.w, both Spectra Physics Excelsior models, and a 638 nm 40 mW laser diode (cat # L638P040, ThorLabs) in a mounting system with temperature and drive current control (cat # LTC56A, ThorLabs), with temperature maintained at 25°C. The lasers were then coupled into a single-mode, polarization-maintaining fiber optic patch cable (P5-405BPM-FC-5, Thorlabs) using an achromatic fiber port (cat # PAF2-A4A, ThorLabs) installed in a fiber port adaptor (cat # CP08FP). The fiber delivered the light into the scanning unit (M612, ISS Inc.). The M612 delivered the excitation light through the right-side port of our microscope using either a dual bandpass/longpass dichroic beamsplitter (ZT488/561rpc, Chroma Technology Corporation) or a quadruple bandpass dichroic beamsplitter (405/488/561/635 nm lasers BrightLine® quad-edge laser dichroic beamsplitter, cat # FL-007046, SemRock). The quadruple dichroic filter was used when the 638 nm laser diode was used. The fluorescence was collected along the same path and focused onto an avalanche photodiode (SPCM-AQRH-44-BR2, Excelitas) (APD). Emission filters were placed in front of the APD detector depending on the laser used: 525/50 nm bandpass (cat # ET525/50m, Chroma Technology) for 488 nm, 570 nm long pass (cat # ET570lp, Chroma Technology) for 561 nm, or 640 nm longpass (cat # BLP01-633R-25, Semrock) for 638 nm. The APD was operated in the photon counting mode, with counts sampled by a USB-connected data acquisition card. Data were acquired by storing the photon counts in time-tagged mode, allowing the counts to be binned at lower frequencies during data analysis. The maximum sampling frequency used was 2 MHz. Data was analyzed on a desktop computer using VistaVision (ISS).

### Photobleaching Data Acquisition and Analysis

The cells were chosen so that, for each power setting, cells of similar intensity were selected for measurement. The PMT gain was varied to place the average intensity in the middle of the detector’s dynamic range (∼1000-2000 a.u., out of a maximum of 4096 a.u.). Fifty frames were obtained by unidirectional line scanning using a commercial laser scanning confocal system (LSM 710, Zeiss). For each fluorophore/power combination, three cells were measured. Data were analyzed using Fiji software (ImageJ) and MATLAB (Mathworks). Frames 3-25 were isolated, and the intensity was normalized to the initial intensity. Frame 3 was considered as t = 0 for this analysis. The data were then fit to the function

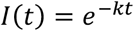

using a nonlinear least squares approach. Rate constants (k) were fit using data from three cells for each fluorophore/power setting, and the average and standard deviation of the rate constants from the three cells are reported in this work, and the relative power *E*_*C*_ is given by

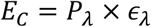

Where *P*_*λ*_ is the incident power at wavelength λ (measured at the objective) and *ϵ*_*λ*_ is the molar extinction coefficient of the FP at wavelength λ. Calculating *E*_*C*_ for each FP used allows us to compare the photostability of FPs that absorb at different wavelengths while accounting for the power output of the laser lines used.

### Calibration of HaloTag Ligand Labeling Efficiency

HEK293T cells were seeded in a 35 mm plastic dish at 15% confluency and incubated for 20 hours, when cells were transfected with 1 µg HaloTag-mEGFP plasmid with 3 µL PEI transfection reagent in 200 µL of FBS-free DMEM medium. Transfected cells were replated in PLL-coated 96-well plates and incubated overnight. Cells were then stained with 200 nM JF646 for different amounts of time (2, 4, 6, 10, and 24 hours). After the indicated time, cells were destained with DPBS containing Ca^2+^/Mg^2+^. Imaging medium was used to incubate cells during experiments. Fluorescence images were collected using the ImageXpress Pico Automated Cell Imaging System (Molecular Devices), which measures the fluorescence intensity of JF646 and mEGFP in each cell at each time point. A 4.8% imaging area and a 3×3 imaging pattern were used for imaging each well with a 10× objective lens. HaloTag-JF646 was imaged using the Cy5 channel (excitation 610-650 nm, emission 675-720 nm), and mEGFP was imaged using the FITC channel (excitation 445-485 nm, emission 509-539 nm). Images were analyzed with ImageJ to quantify the labeling efficiency. For each cell, the fluorescence intensity ratio was calculated by (integrated density of HaloTag-JF646) / (integrated density of mEGFP). The average ratio from four technical replicates was then calculated for each well and plotted against staining time using MATLAB, with the maximum value normalized to 1.0.

### Measurement of APD Dead Time

The relationship between the observed fluorescence intensity, variance, and detector parameters for a constant light source are given by (52)

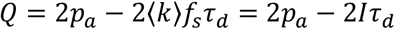

Where *p*_*a*_ is the afterpulsing probability, ⟨k⟩ is the average photon count rate in counts per sample period, *f*_*s*_ is the sampling frequency in Hz, *τ*_*d*_ is the detector dead time in seconds, and *I* is the fluorescence intensity in counts per second, equivalent to ⟨k⟩ × *f*_*s*_. Mandel’s Q parameter is given by

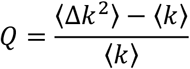

Where ⟨k⟩ and ⟨Δk^2^⟩ are the average and variance of the fluorescence intensity. To measure the afterpulsing probability and dead time, we measured Mandel’s Q parameter for a solution of highly concentrated dye solution (20 µM of Atto565) using a 561 nm laser line. At this concentration, fluctuations due to molecular diffusion are absent, and the only fluctuations present arise from the detector’s response. To generate a linear fit to the model, we increased the fluorescence intensity by increasing the power of the excitation laser. We calculated Q directly from the fluorescence intensity traces and fit the data to a linear model Q = 2*p*_*a*_ − 2*Iτ*_*d*_. Since correcting for afterpulsing effects has little effect on the resulting PCH fits(52), we only include the dead time in our fitting procedure.

### Measurement of FFS Observation Volume

We related the number of particles recovered by PCH to the absolute concentration by independently measuring the observation volume of our microscope using a solution of fluorescent dye with known concentration (Figure S1). We prepared a 20 nM solution of Atto655 (free acid) diluted in MilliQ purified water with 0.05% Tween-20 (v/v). The detergent was included to reduce the adsorption of dye molecules to the surface of the coverslip. The concentration of the dye stock solution was measured independently in the detergent/water solution using absorption spectroscopy with a Cary 3500 UV-Vis Compact Peltier Spectrophotometer (Agilent) and an extinction coefficient of ε_655_ = 125,000 *M*^−1^*cm*^−1^. The dye was diluted to 20 nM in a single step. A 400 µL aliquot of the diluted dye solution was added to the same glass-bottom dishes used for cell experiments. The dye was excited using the 638 nm laser line, and the laser power was adjusted until approximately 2 µW/cm^2^ power was observed when measured just before the objective lens. We measured laser power by using a threaded adaptor (cat # SM1BC, ThorLabs) that connects our power meter probe (cat # S121C, ThorLabs) to the microscope turret, allowing for precise and consistent power measurements each day. Fluorescence was collected with a 640 nm longpass emission filter. Single-point FFS data was collected for 30 seconds. PCH data was binned at 20 kHz and fit by fixing the detector dead time to 15 ns and sampling frequency at 20 kHz in the fitting algorithm. For this measurement, the pinhole was set to approximately 1 Airy unit (56 µm for our system with 80× total magnification). All measurements and calculations in this work that refer to absolute concentrations were measured using this pinhole setting.

Fitting the resulting histogram using the PCH model resulted in a recovered number of particles N. To calculate the observation volume, we applied the relation

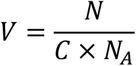

Where V is the observation volume in liters, N is the number of particles measured by PCH, C is the known concentration of the dye solution in molar units, and *N*_*A*_ is Avogadro’s constant.

We adjusted the laser power until PCH measurements confirmed consistent PCH fits. We calibrated the system each day to account for any changes in the microscope’s collection efficiency function (slight changes in the size of the variable pinhole, temperature effects on alignment, etc.) that might occur from day to day.

### Fluorescence Staining of Cells for FFS

For all FCS/PCH experiments, HaloTag fusion constructs were labeled with 200 nM Janelia Fluor 646 HaloTag ligand (Promega, catalog # HT1060) in growth medium for at least 6 hours, followed by two destaining washes with DPBS containing Ca^2+^/Mg^2+^. Imaging medium supplemented with 25 mM HEPES, pH 7.4 was used during imaging experiments, which occurred at ambient temperature (22-23°C).

### PCH of Live Cells

All PCH measurements were made using our custom FFS microscope. Fluorescence intensity was measured for either 10 seconds at each spot (fluorescent protein fusion constructs) or 30 seconds at each spot (HaloTag fusion constructs), with at least 6 spots measured for each cell. We fit the binned counts from each spot (20 kHz binning frequency, 15 ns dead time) using PCH. Each trajectory was inspected manually, and trajectories that included significant artifacts due to cell movement were not analyzed. When measuring HaloTag fusion constructs, small intensity drifts (<10% change over the entire trajectory) were occasionally identified, and analysis was restricted to ≥ 10-second regions where no intensity drift was observed. The brightness of each cell was calculated as the average of the brightness values from each spot to calculate the average molecular brightness. Any individual spot with a brightness that differed from the average by more than 1.5σ, where σ represents the sample standard deviation, was excluded in the final analysis.

For light activation, the entire dish was exposed to ∼ 6 mW/cm^2^ of blue light for 5 minutes. Power density was measured using a power meter (cat # PM100D, ThorLabs). Immediately after turning off the blue light, a single measurement of 30 seconds was taken for each cell to minimize the effect of dark state recovery during the measurement.

### Calculation of Protomer Concentration

For light-exposed cells, the HaloTag fusion constructs are expected to exist as a population of mixed oligomers. To calculate the protomer concentration in cells, we used the following equation

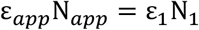

Thus, one can calculate a “protomer equivalent” concentration for a sample of arbitrary brightness by dividing the total count rate by the monomeric brightness. After performing this calculation, we convert it to absolute concentration using the calibrated measurement of the observation volume (1.02 fL). Finally, we divide by the labeling efficiency (70%) to account for unlabeled molecules. Our protomer concentration, therefore, represents the expected concentration of all (labeled + unlabeled) protomers in the cell at the measurement location. These corrections were made before adjusting the molecular brightness into relative units.

### Calculation of Corrected Relative Brightness Values

For all measurements reported in **Figures 3-4**, we normalized the molecular brightness values to the monomeric standard and corrected the relative molecular brightness values to account for incomplete labeling, thereby simplifying the interpretation of the scatter plots. We used two equations to correct the measured brightness ε_*app*_. First, we normalize the brightness to the monomeric brightness:

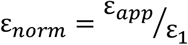

If ε_*app*_ ≤ ε_1_ (due to experimental uncertainty or noise), we use ε_*norm*_ directly as the corrected relative brightness. If ε_*app*_ > ε_1_, then we use the following correction (34)

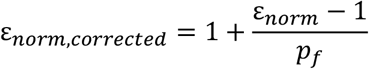

Where *p*_*f*_ is the fluorescence probability determined by our calibration measurement of the HaloTag-JF646 tandem dimer and monomer (0.7 for this work). After these corrections, a relative brightness value of 2 corresponds to the relative brightness expected for a constitutive dimer. This can be seen by substituting our measured (uncorrected) brightness ratio of ε_*norm*_ = 1.7 for the HaloTag-JF646 dimer:monomer, and the corresponding fluorescence probability *p*_*f*_ = 0.7:

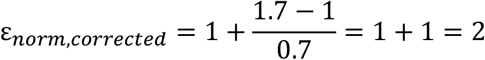

### Light-induced Lytic Cell Death

Optogenetic activation was carried out using a custom-built blue LED system (465 nm, 0.5 mW/cm^2^, 15 s on/off cycle) for 12 hours. Plasma membrane rupture was assessed by staining cells with 250 nM SytoxGreen, and fluorescence imaging was performed using a Leica DMI8 fluorescence microscope with a 10X objective lens to visualize SytoxGreen (GFP channel, excitation 450-490 nm, emission 500-550 nm) and transgene expression (TXR channel, excitation 540-580 nm, emission 592-668 nm).

## Supporting information

SupportingInformation

## Acknowledgments

This work is supported by the American Parkinson Disease Postdoctoral Fellowship (Award No. 978673) (T.C.). National Institute of General Medical Sciences (R01GM132438) and National Institute of Mental Health (R01MH124827), NSF (Award No. 2121003), and an NSF Science and Technology Center for Quantitative Cell Biology Grant (Award No. 2243257) (K.Z.). We are grateful to Dr. Yuansheng Sun from ISS for insightful comments and support in building the microscope and PCH analysis.

## Notes

### Competing Interest Statement

The authors have declared no competing interest.

