## SupportingInformation for "Resolving Oligomeric States of Photoactivatable Proteins in Live Cells via Photon Counting Histogram Analysis"

**This PDF file includes:**

Supporting text
Figures S1 to S3
Tables S1 to S2

### Supplementary Materials and Methods

#### Reagents

The following plasmids were ordered from Addgene: pCaggs-mGold (#157996), pHR-hSyn-EGFP (#114215), pmScarlet_C1 (#85042), mCherry-CRY2clust (#105624). HaloTag was a kind gift from Prof. Jason D. Shephard's lab at the University of Utah. FastDigest BamHI (cat # FD0054), BglII (cat # FD0083), KpnI (cat # FD0524), HindIII (cat # FD0504), and NotI (cat # FD0593) were purchased from ThermoFisher Scientific. rCutSmart buffer (cat # B6004S), T4 DNA ligase buffer (cat # B0202S), recombinant shrimp alkaline phosphatase (rSAP, cat # M0371S), T4 DNA ligase (cat # M0202S), and Q5 Site-Directed Mutagenesis Kit (cat # E0554S) were purchased from New England Biolabs. InFusion HD Cloning Kit (cat # 639650) was purchased from Takara Bio USA, Inc. Atto655 (free acid, cat # AD 655-21) and Atto565 (free acid, cat # AD 565-21) were purchased from ATTO-TEC GmbH. JaneliaFluor 646 HaloTag ligand was purchased from Promega (cat # HT1060). Dulbecco's modified eagle medium (DMEM) was purchased from Corning (cat # 10-013-CV). Opti-MEM with L-glutamine and no phenol red was purchased from Gibco (cat # 11058021). Penicillin/streptomycin (pen/strep) was purchased from Corning (cat # 30-001-CI). Fetal bovine serum was purchased from Avantor (cat # 76419-584). PEI Max transfection reagent was purchased from Polysciences (cat # 24765(1)). Glass-bottom 12-well plates (cat # P12-1.5H-N) and 96-well plates (cat # P96-1.5H-N) were purchased from Cellvis. Glass-bottom 35 mm dishes were purchased from Mattek (cat # P35G-1.5-14-C). Poly-L-lysine was purchased from Millipore Sigma (cat # P1274). Trypsin was purchased from Gibco (cat # 25200-056).

#### Plasmid Construction

Plasmids were generated by InFusion assembly or restriction digest. Plasmids containing only monomeric fluorescent proteins/HaloTag were generated using InFusion assembly. The pCS2+ vector backbone with cytomegalovirus (CMV) promoter was linearized by digesting with BamHI, and the inserts were amplified with overlap extension PCR to generate complementary overhangs. The insert and vector were combined with InFusion HD Enzyme premix contained within the InFusion HD Cloning Kit. The monomer plasmids contained a BamHI before the Kozak sequence, a BglII site after the coding sequence but before the stop codon, and a KpnI site after the stop codon. To generate plasmids containing tandem oligomers, the monomer plasmids were digested with either BglII/KpnI (to generate the new vector backbone) or BamHI/KpnI (to generate the new insert). To generate tandem oligomer plasmids with the flexible linker glycine/serine linker, the second gene sequence was amplified with PCR using the monomeric plasmid as a template, with the linker and BamHI restriction site added via primer extension. The PCR product was then digested with BamHI/KpnI. The monomer was digested with BglII/KpnI and ligated with the digested PCR product. Phosphates were removed from each vector with rSAP prior to ligation. T4 DNA ligase was used for all ligation reactions. Fusions of HaloTag with VfAuLOV and AtCRY2 variants were assembled with restriction digest cloning using templates that had previously been generated using InFusion cloning. The vector plasmid contained the HaloTag sequence followed by the flexible linker/EcoRI site before the stop codon, and a KpnI site after the stop codon. The insert plasmids contained an N-terminal EcoRI site followed by either the LOV or CRY2 sequences, a stop codon, and a KpnI site. Plasmids were double digested with EcoRI and KpnI and ligated. Colonies were screened with restriction digest or colony PCR, and the final product was confirmed by sequencing. For longer, repetitive sequences such as tandem oligomers, we used Plasmidsaurus sequencing service.

For FFS experiments, we often found that the CMV promoter produced very high protein concentrations that are not compatible with FFS. For this reason, we generated plasmids with the Ubiquitin C promoter controlling gene expression of our constructs, which expresses proteins constitutively but at lower levels. These plasmids were generated by digesting the pCS2-CMV plasmids with HindIII/NotI (which are placed before and after the gene sequence) to generate the insert. The insert was ligated into the vector, which was generated by digesting UBC-EGFP (Addgene #169746) with HindIII/NotI. The cloning proceeded as above for other ligation reactions. Site-directed mutagenesis was further performed on the final HaloTag-VfAuLOV/AtCRY2 plasmids to remove trailing residues on the C-terminus (left over by restriction digestion used to build the templates). The final constructs for all plasmids containing HaloTag are in the process of depositing with Addgene. Information of plasmids and primers used in this work is shown in **Table S1** and **S2**, respectively.

#### Cell Culture and Transfection

Human embryonic kidney cells (HEK293T, ATCC #CRL-3216) were used for all experiments. Cells were maintained in growth medium (DMEM supplemented with 10% FBS and 100 U/mL penicillin, 100ug/mL streptomycin). Cells were passaged by trypsin digestion and reseeded at 10-40% density. For imaging experiments, cells were seeded in glass-bottom (no. 1.5 coverslip) 12-well tissue culture plates (Cellvis) or glass-bottom 35 mm dishes with a 14-mm coverslip diameter (Mattek). The plates were coated with poly-L-lysine for at least 30 minutes prior to seeding cells. For light-induced cell death, HEK293T cells were seeded in a 48-well plate with 300 µL of Dulbecco's Modified Eagle Medium (DMEM) supplemented with 10% fetal bovine serum (FBS) and penicillin-streptomycin. Cells were incubated at 37°C with 5% $CO_2$ for 24 hours before transfection.

Cells were transfected at 40-60% confluence with PEI Max using 0.5-1 µg of DNA per well/dish. In some experiments, protein expression levels were lowered further by diluting plasmid DNA containing our constructs with empty pCS2+ plasmid DNA. After optimization, we found that 50 ng of construct DNA + 450 ng of empty pCS2+ plasmid DNA provided optimal expression levels without reducing transfection efficiency too much. PEI:DNA ratio was kept at 3:1 (v/w) and mixed with 60 µL (when using 35 mm dishes) or 100 µL (when using glass-bottom 12-well plates) of DMEM per well. After thoroughly mixing the transfection solution and incubating at room temperature for at least 15 minutes, the solution was added to cells drop by drop. Cells were placed in the incubator for 4 hr, after which the media was changed. Before imaging, growth media was replaced with imaging media. Imaging media was prepared by mixing Opti-MEM with L-glutamine (cat # 11058021, Gibco) with 10% fetal bovine serum and antibiotics as above. Imaging media contained no phenol red. When cells remained out of the incubator for several hours for imaging, 25 mM HEPES, pH 7.4 (cat # 25-060-CI from Corning) was added to the imaging media. Cells were kept at ambient temperature for no more than 3 hours. For light-induced cell death, 0.3 µg of optogenetic RIPK3-encoding plasmid DNA was complexed with 0.9 µg of PEI MAX in serum-free DMEM to a final volume of 30 µL. The transfection mixture was added dropwise to the center of each well and incubated for 4 hours, after which the medium was replaced with complete DMEM to minimize cytotoxicity. At 20 hours post-transfection, the medium was replaced again to remove spontaneously dead cells while preserving viable cells.

#### Fluorescence Confocal Imaging

Data presented in Figure 1A-B was collected using an LSM 710 laser scanning confocal microscope (Zeiss). All images were obtained using the 40×, 1.2 NA water immersion objective.

The excitation was either a 488 nm, 514 nm, or 561 nm laser line. Power measurements were taken at the objective for different settings of the acousto-optic tunable filter that adjusts the laser power on our system. Images were acquired using a pixel dwell time of 12.54 microseconds, 256 × 256 pixels per image, with a pixel size of 100 nm. The pinhole size was always set to 1 Airy unit.

**Structural Model Analysis**

Molecular graphics and analyses shown in Figure 4 were performed with UCSF ChimeraX, developed by the Resource for Biocomputing, Visualization, and Informatics at the University of California, San Francisco, with support from National Institutes of Health R01-GM129325 and the Office of Cyber Infrastructure and Computational Biology, National Institute of Allergy and Infectious Diseases.

Structural alignment of AtCRY2 and MmCRY2 was done using the "Matchmaker" tool in ChimeraX using $C^{\alpha}$ atoms. The Needleman-Wunsch alignment algorithm was used with the BLOSUM-62 substitution matrix. Secondary structure weighting was set to 0.3. The intra-helix gap opening penalty and the intra-strand gap opening penalty were set to 18. All other gaps were assigned a penalty of 6. The resulting amino acid sequence alignment was displayed in Figure S2 using Jalview and was used to identify residues on MmCRY2 that correspond to the HT interface residues of AtCRY2.

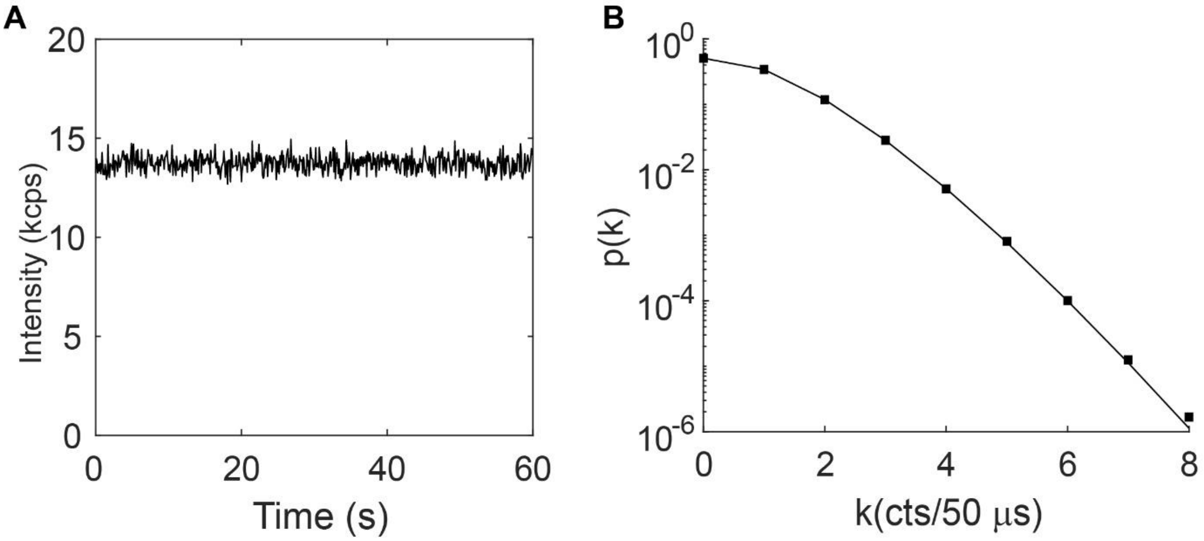


**Figure S1. Representative calibration standard for FCS measurements.** A 20 nM solution of Atto655 in Milli-Q water + 0.05% Tween-20 (v/v) was measured at a single position within the sample above the coverslip (A) and fit to the PCH model (B). PCH data is shown as black squares with the fit to the model drawn as a black line. For this sample, the molecular brightness was 1,200 counts per second per molecule (cpsm), the number of particles was 12, and the chi-square value was 0.69.

```
AtCRY2     1 ---------------MKMD-KKTIVWFRRDLRIEDNPALAAAAHEG-SVFPVF
MmCRY2     1 MAAAAVVAATVPAQSMGADGASSVHWFRKGLRLHDNPALLAAVRGARCVRCVY

AtCRY2    37 IWCPEEEGQFYPGRASRWWMKQSLAHLSQSLKALGSDLTLIQTHNTISAILDC
MmCRY2    54 ILDPWFAASSSVGINRWRFLLQSLEDLDTSLRKLNSRLFVVR-GQPADVFPRL

AtCRY2    90 IRVTGPTKVVFNHLYDPVSLVRDHTVKEKLVERGISVQSYNGDLLYEPWEIYC
MmCRY2   106 FKEWGVTRLTFEYDSEPFGKERDAAIMKMAKEAGVEVVTENSHTLYDLDRIIE

AtCRY2   143 EKG-KPFTSFNSYWKKCLDMSI-ESVMLPPPWRLMPITAAA------EAIWAC
MmCRY2   159 LNGQKPPLTYKRFQALISRMELPKKPAVAVSSQQMESCRAEIQENHDDTYGVP

AtCRY2   188 SIEELGLENEAEKPSNALLTRAWSPGWSNADKLLNEFIEKQLIDYAKNSKKVV
MmCRY2   212 SLEELGFPTEGLGPA------VWQGGETEALARLDKHLERKA--WVANYERPR

AtCRY2   241 GN------STSLLSPYLHFGEISVRHVFQCARMKQIIWARDKNSEGEESADLF
MmCRY2   257 MNANSLLASPTGLSPYLRFGCLS--CRLFYYRLWDLYKKVKRNS---TPPLSL

AtCRY2   288 LRGIGLREYSRYICFNFP-FTHEQSLLSHLRFFPWDADVDKFKAWRQGRTGYP
MmCRY2   305 FGQLLWREFFYTAATNNPRFDRMEGNPICIQ-IPWDRNPEALAKWAEGKTGFP

AtCRY2   340 LVDAGMRELWATGWMHNRIRVIVSSFGVKF-LLLPWKWGMKYFWDTLLDADLE
MmCRY2   357 WIDAIMTQLRQEGWIHHLARHAVACFLTRGDLWVSWESGVRVFDELLLDADFS

AtCRY2   392 CDILGWQYISGSIPDGHELDRLDNPALQGAKYDPEGEYIRQWLPELARLPTEW
MmCRY2   410 VNAGSWMWLSCS-AFFQQFFHCYCPVGFGRRTDPSGDYIRRYLPKLKGFPSRY

AtCRY2   445 IHHPWDAPLTVLKASGVELGTNYAKPIVD-IDTARELLAK
MmCRY2   462 IYEPWNAPESVQKAAKCIIGVDYPRPIVNHAETSRLNIER
```

**Figure S2. Structural alignment of AtCRY2 and MmCRY2.** The dark-state monomer of AtCRY2 (PDB 6K8I) was aligned to the structure of FAD-bound MmCRY2 (PDB 4I6G) using the "Matchmaker" tool of ChimeraX. The alignment was used to highlight residues of the HT interface in the AtCRY2 and their matching residues in MmCRY2.

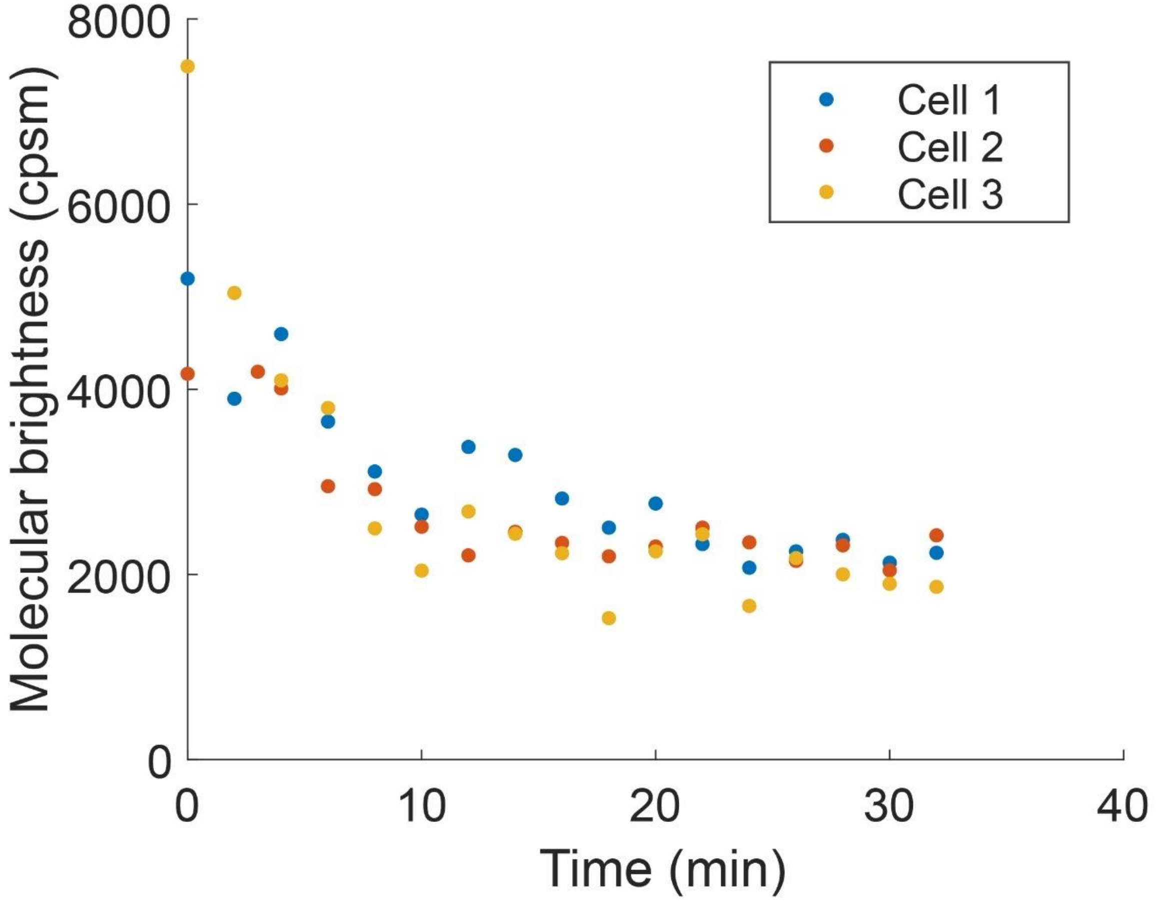


**Figure S3. Dark state recovery of AtCRY2 in live HEK293T cells.** After a 5-minute blue light exposure, the molecular brightness was measured at the indicated times after the blue light was turned off. Each measurement lasted 30 seconds. Each color (orange, yellow, blue) represents a separate cell.

**Table S1:** Plasmids used in this work.

| Name | Source |
|---|---|
| pUBC-1X-HaloTag | This study |
| pUBC-2X-HaloTag | This study |
| pCMV-1X-mEGFP | This study |
| pCMV-2X-mEGFP | This study |
| pCMV-1X-mCherry | This study |
| pCMV-2X-mCherry | This study |
| pCMV-1X-mScarlet | This study |
| pCMV-1X-mGold | This study |
| pUBC-HaloTag-AuLOV | This study |
| pUBC-HaloTag-bZipAuLOV | This study |
| pUBC-HaloTag-AtCRY2 | This study |
| pUBC-HaloTag-AtCRY2(W374A) | This study |
| pUBC-HaloTag-HsCRY2 | This study |
| mCh-RIPK3-AuLOV | Reference 41 in main text |
| mCh-RIPK3-bZipAuLOV | Reference 41 in main text |
| mCh-RIPK3-AtCRY2 | Reference 41 in main text |

**Table S2:** Primers used in this work.

| S. No. | Name | Sequence | Purpose |
|---|---|---|---|
| 1 | mEGFP.FOR | CACCCAGTCCaagCTGAGCAAAGAC | A206K mutation to build monomeric EGFP |
| 2 | mEGFP.REV | CTCAGGTAGTGGTTGTCG | A206K mutation to build monomeric EGFP |
| 3 | 1X-FP.FOR | TCTTTTTGCAGGATCCgccaccatggtgagc | Build fluorescent protein monomer plasmids |
| 4 | 1X-FP.REV | CGAATCGATGGGATCTTAAGATCTcttgtacagctcgtccatgcc | Build fluorescent protein monomer plasmids |
| 5 | 1X-HT.FOR | TCTTTTTGCAGGATCCgccaccatgGCAGAAATCGGTACTGGC | Build HaloTag monomer plasmid |
| 6 | 1X-HT.REV | CGAATCGATGGGATCttaagatctAGTGGTTGGCTCGCC | Build HaloTag monomer plasmid |
| 7 | HT-Linker.FOR | ATGATCGGATCCGGAGGTGGAAGCGGTGGAGGCTCAGCAGAAATCGGTACTGGCTTTCC | Add flexible linker to HaloTag tandem dimer |
| 8 | HsCRY-Insert.FOR | GGAGGCTCAGAATTCatggcggcaactgtggca | Amplify Human CRY2 PHR insert |
| 9 | HsCRY2-Insert.REV | AGGCCTTGAATTtcaccgtgggtagtccacacca | Amplify Human CRY2 PHR insert |
| 10 | HsCRY2-Vector.FOR | tgaAATTCAAGGCCTCTCGAGC | Amplify pUBC-HaloTag plasmid vector |
| 11 | HsCRY2-Vector.REV | GAATTCTGAGCCTCCACCGC | Amplify pUBC-HaloTag plasmid vector |
| 12 | RemoveRS.FOR | TGAAATTCAAGGCCTCTC | Remove leftover restriction enzyme recognition sequence |
| 13 | RemoveRS-AuLOV.REV | TTTGCGTCTCAGCATATTG | Remove restriction sequence from LOV-based plasmids |
| 14 | RemoveRS-CRY2.REV | ggcagcaccgatcataatc | Remove restriction sequence from CRY2 plasmids |
| 15 | Nterm-HT-Linker.FOR | ggtggaggctcaGAATTCAAGGCCTCTCGAG | Add flexible linker to HaloTag |
| 16 | Nterm-HT-Linker.REV | gcttccacctccAGTGGTTGGCTCGCC | Add flexible linker to HaloTag |
| 17 | Cterm-AuLOV.FOR | CCATCGATTCGAATTCGCCCAGAGCTCACCAGAG | Generate AuLOV template with EcoRI site |
| 18 | Cterm-bZipAuLOV.FOR | CCATCGATTCGAATTCGGTAGTATAAGCTCAGAACTTACTGAAG | Generate bZipAuLOV template with EcoRI site |
| 19 | Cterm-LOV.REV | GAGAGGCCTTGAATTTTAGCTAGCTTTGCGTCTCAGCATATTGGAGG | Generate LOV sequences with EcoRI site |
| 20 | Cterm-CRY2.FOR | CCATCGATTCGAATTCatgaagatggacaaaaagaccatcgt | Generate CRY2 sequence with EcoRI site |
| 21 | Cterm-CRY2.REV | GAGAGGCCTTGAATTtcagctagcggcagcaccgatcataat | Generate CRY2 sequence with EcoRI site |
